# Design of Epitope Based Peptide Vaccine Against Pseudomonas Aeruginosa Fructose Bisphosphate Aldolase Protein using Immunoinformatics

**DOI:** 10.1101/720730

**Authors:** Mustafa Elhag, Ruaa Mohamed Alaagib, Nagla Mohamed Ahmed, Mustafa Abubaker, Esraa Musa Haroun, Sahar Obi, Mohammed A. Hassan

## Abstract

*Pseudomonas aeruginosa* is common pathogen that is responsible of serious illnesses hospital acquired infection as ventilator associated pneumonia and various sepsis syndrome. Also it is a multidrug resistant pathogen recognized for its ubiquity, its intrinsically advanced antibiotic resistant mechanisms. generally affects the immuonocompromised but can also infect the immunocompetent as in hot tub folliculitis. There is no vaccine against it available till now. This study predicts an effective epitope-based vaccine against Fructose bisphosphate aladolase (FBA) of *Pseudomonas aeruginosa* using immunoinformatics tools. The sequences were obtained from NCBI and prediction tests took place to analyze possible epitopes for B and T cells. Three B cell epitopes passed the antigenicity, accessibility and hydrophilicity tests. Six MHC I epitopes were the most promising, while four from MHC II. Nineteen epitopes were shared between MHC I and II. For the population coverage, the epitopes covered 95.62% of the alleles worldwide excluding certain MHC II alleles. We recommend invivo and invitro studies to prove it’s effectiveness.

## INTRODUCTION

*Pseudomonas aeruginosa* is motile, non-fermenting, gram negative opportunistic bacterium that implicated in respiratory infections, urinary tract infections, gastrointestinal infections, keratitis, otitis media, and bacteremia in patients with compromised host defences (e.g., cancer, burn, HIV, and cystic fibrosis). [1] Intensive care units (ICU) hospitalized patients constitute one of the risk group that are more susceptible of acquiring pseudomonas infections as they may develop ventilator-associated pneumonia (VAP) and sepsis.[2–4] This organism is a ubiquitous and metabolically versatile microbe that flourishes in many environments and possesses many virulence factors that contribute to its pathogenesis [1] According to data from Centers for Disease Control, *P. aeruginosa* is responsible for millions of infections each year in the community, 10–15% of all healthcare-associated infections, with more than 300,000 cases annually in the EU, USA and Japan. [5] It is a common nosocomial pathogen, [6, 7] that causes infections with a high mortality rate [8, 9] which is attributable to that the organism possesses an intrinsic resistance to many antimicrobial agents., [10] and the development of increased, multidrug resistance in healthcare settings, [11–13] both of which complicate anti-pseudomonal chemotherapy. As a result, it remains difficult to combat *P.aeruginosa* infections despite supportive treatments. Vaccines could be an alternative strategy to control *P.aeruginosa* infections and even reduce antibiotic resistance; however no *P.aeruginosa* vaccine is currently available. [14] Doring and pier (2008) represented that the serious obstacle to the development of a globally effective anti–P. aeruginosa vaccine are due to antigenic variability of microorganism that enable it to easily adapt to different growth condition and escapes host immune recognition, and to the high variability of the proteins among different *P. aeruginosa* strains and within the same strain, grown in diverse environmental conditions. [15]

So far *P.aeruginosa* vaccine candidate have been found by classical approach. Integrated genomics and proteomics approaches have been recently used to predict vaccine candidates against *P. aeruginosa*. [16] Although several vaccine formulations have been tested clinically, none has been licensed. [15, 17] The search for new targets or vaccine candidates is of high paramount. Bioinformatics-based approach is a novel platform to identify drug targets and vaccines candidates in human pathogens. [18, 19] Thus the present study aimed to design effective peptide vaccine against *P.aeruginosa* using computational approach through prediction of highly conserved T and B cell epitopes from the most conserved and highly immunogenic protein Fructose Bisphosphate Adolase (FBA) protein. This is the first study that predicts epitope-based vaccine from the candidates moonlighting protein against *P.aeruginosa*. This technique has been successfully used by several authors to identify drug target vaccine candidates. These type of vaccines are easy to produce, specific, capable to keep away from undesirable immune responses, reasonably and also safe when compared to the usual vaccines like killed vaccines and attenuated vaccine. [20]

## MATERIALS AND METHODS

### Protein sequence retrieval

A total of 20,201 strains of *Pseudomona Aeruginosa* FBA were retrieved in FASTA format from National Center for Biotechnology Information (NCBI) database (https://ncbi.nlm.nih.gov) on May 2019.The protein sequence had length of 354 with name fructose-1,6-bisphosphate aldolase.

### Determination of conserved regions

The retrieved sequences of *Pseudomona Aeruginosa* FBA were subject to multiple sequence alignment (MSA) using ClustalW tool of BioEdit Sequence Alignment Editor Software version 7.2.5 to determine the conserved regions. Also molecular weight and amino acid composition of the protein were obtained.[21, 22]

### Sequenced-based method

The reference sequence (NP_249246.1) of *Pseudomona Aeruginosa* FBA was submitted to different prediction tools at the immune epitope database (IEDB) analysis resource (http://www.iedb.org/) to predict various B and T cell epitopes. Conserved epitopes would be considered as candidate epitopes for B and T cell. [23]

### B cell Epitope prediction

B cell epitope is the portion of the vaccine that interacts with b lymphocyte which are a type of white blood cell of the lymphocyte subtype. Candidate epitopes were analysed using several B cell prediction methods from IEDB (http://tools.iedb.org/bcell/), to identify the surface accessibility, antigenicity and hydrophilicity with the aid of random forest algorithm, a form of unsupervised learning. The Bepipred Linear prediction 2 was used to predict linear B cell epitope with default threshold value 0.533 (http://tools.iedb.org/bcell/result/). The Emini surface accessibility prediction tool was used to detect the surface accessibility with default threshold value 1.00 (http://tools.iedb.org/bcell/result/). The Kolaskar and Tongaonker Antigenicity method was used to identify the antigenicity sites of candidate epitope with default threshold value 1.032 (http://tools.iedb.org/bcell/result/). The Parker hydrophilicity prediction tool was used to identify the hydrophilic, accessible, or mobile regions with default the threshold value 1.695.[24–28]

### T cell epitope prediction MHC class I binding

T cell epitope is the portion of the vaccine that interacts with T lymphocytes. Analysis of peptide binding to the MHC (Major Histocompatibility Complex) class I molecule was assessed by the IEDB MHC I prediction tool (http://tools.iedb.org/mhci/) to predict cytotoxic T cell epitopes (also known as CD8+ cell). The presentation of peptide complex to T lynphocyte undergoes several steps. Artificial Neural Network (ANN) 4.0prediction method was used to predict the binding affinity. Before the prediction, all human allele length were selected and set to 9 amino acids. The half-maximal inhibitory concentration (IC50) value required for all conserved epitopes to bind at score less than 500 were selected.[29–35]

### T cell epitope prediction MHC class II binding

Prediction of T cell epitopes interacting with MHC Class II was assessed by the IEDB MHC II prediction tool (http://tools.iedb.org/mhcii/) for helper T cell. Which known as CD4+ cell also. Human allele references set were used to determine the interaction potentials of T cell epitopes and MHC Class II allele (HLA DR, DP and DQ). NN-align method was used to predict the binding affinity. IC50 values at score less than 100 were selected. [36–39]

### Population coverage

The population coverage link was selected to analyse the epitopes in IEDB. This tool calculates the fraction of individuals predicted to respond to a given set of epitopes with known MHC restriction (http://tools.iedb.org/population/iedbinput). The appropriate checkbox for calculation was checked based on MHC I, MHC II separately and combination of both.[40]

### Homology Modelling

The 3D structure was obtained using raptorX (http://raptorx.uchicago.edu) i.e a protein structure prediction server developed by Xu group, excelling at prediction 3D structure for protein sequences without close homologs in the protein data bank (PDB). USCF chimera (version 1.8) was the program used for visualization and analysis of molecular structure of the promising epitopes (http://www.cgl.uscf.edu/chimera). [41, 42]

## RESULTS

### B-cell epitope prediction

The reference sequence of Fructose 1,6-Bisphosphate Aldolase was subjected to Bepipred linear epitope prediction, Emini surface accessibility, Kolaskar and Tongaonkar antigenicity and Parker Hydrophilicity methods in IEDB to test for various immunogenicity parameters. Table 1 and figures 1-4.

**Table 1:**
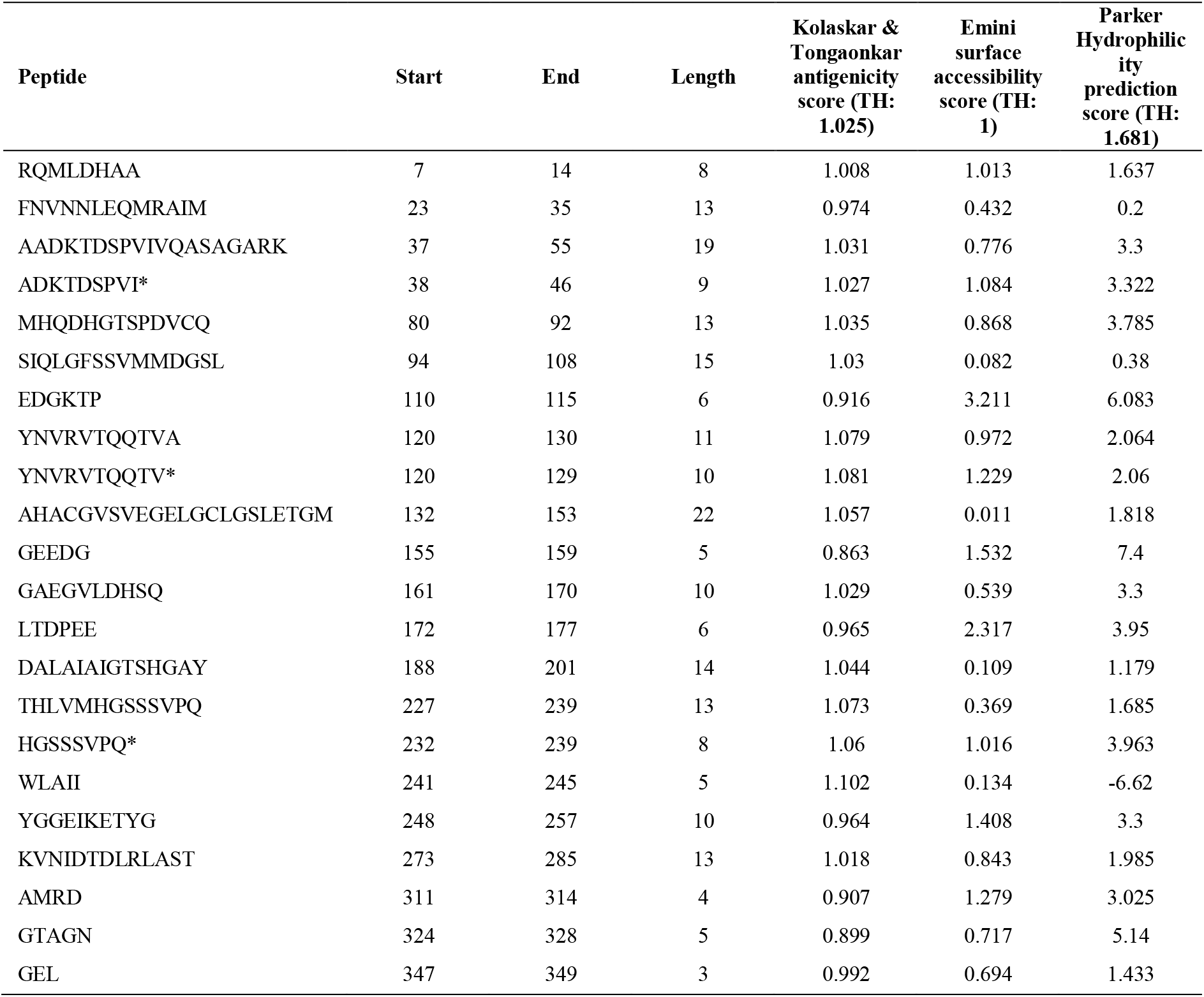
List of conserved peptides with their antigenicity, EMINI surface accessibility and Parker hydrophilicity scores. (*Peptides that successfully passed the three tests).

**Figure 1:**
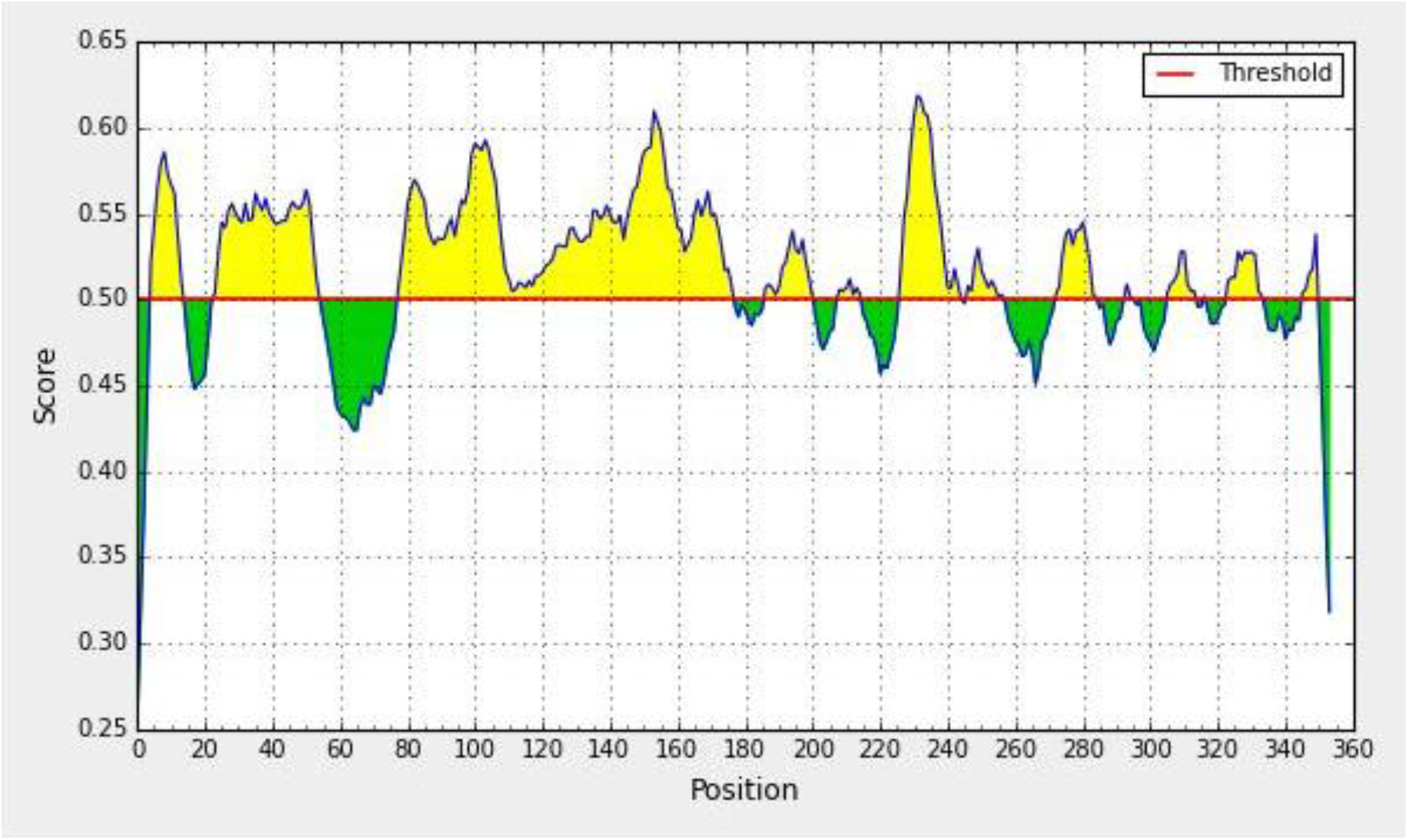
Bepipred Linear Epitope Prediction; Yellow areas above threshold (red line) are proposed to be a part of B cell epitopes and the green areas are not.

**Figure 2:**
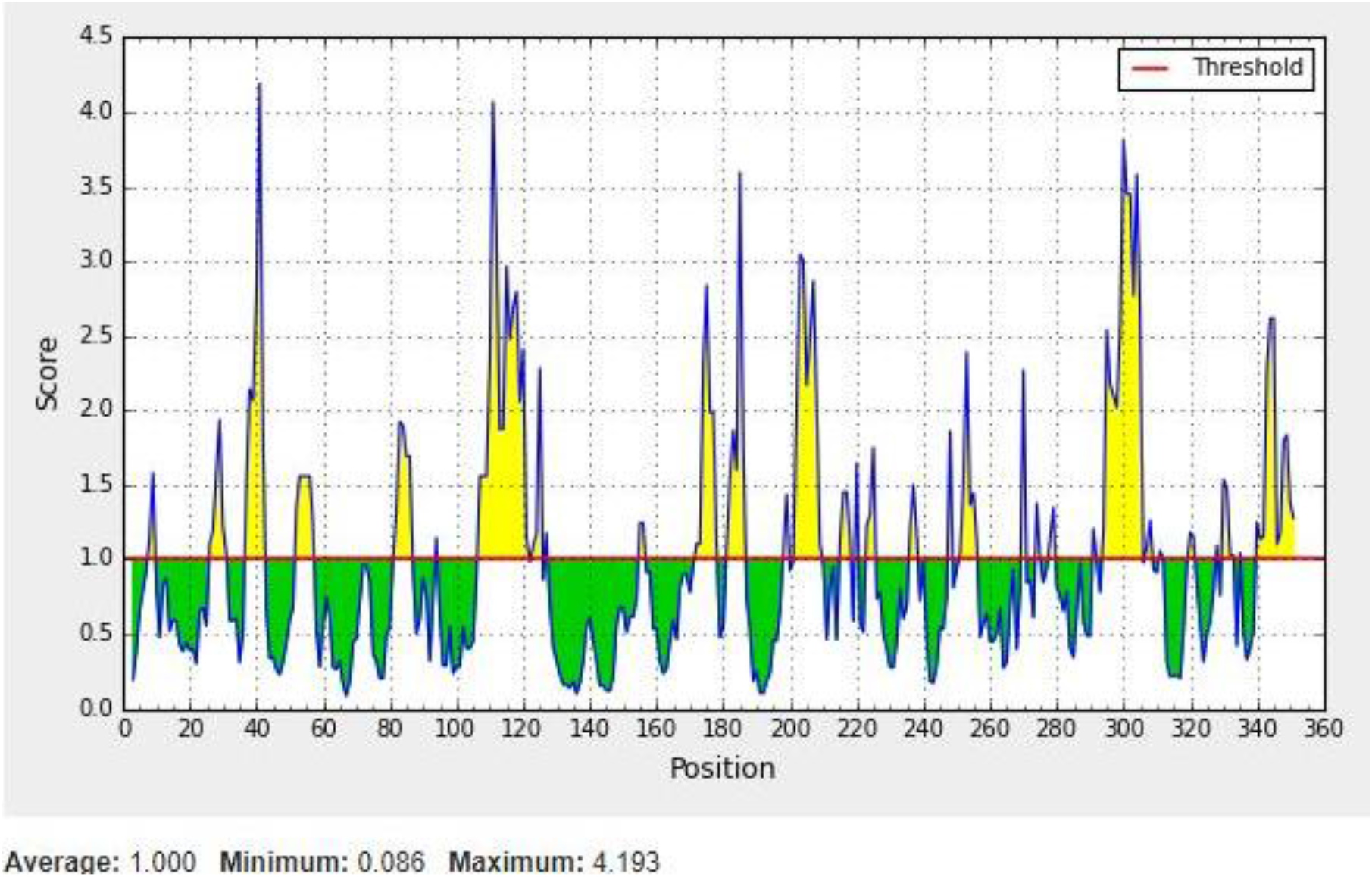
EMINI surface accessibility prediction; Yellow areas above the threshold (red line) are proposed to be a part of B cell epitopes and the green areas are not.

**Figure 3:**
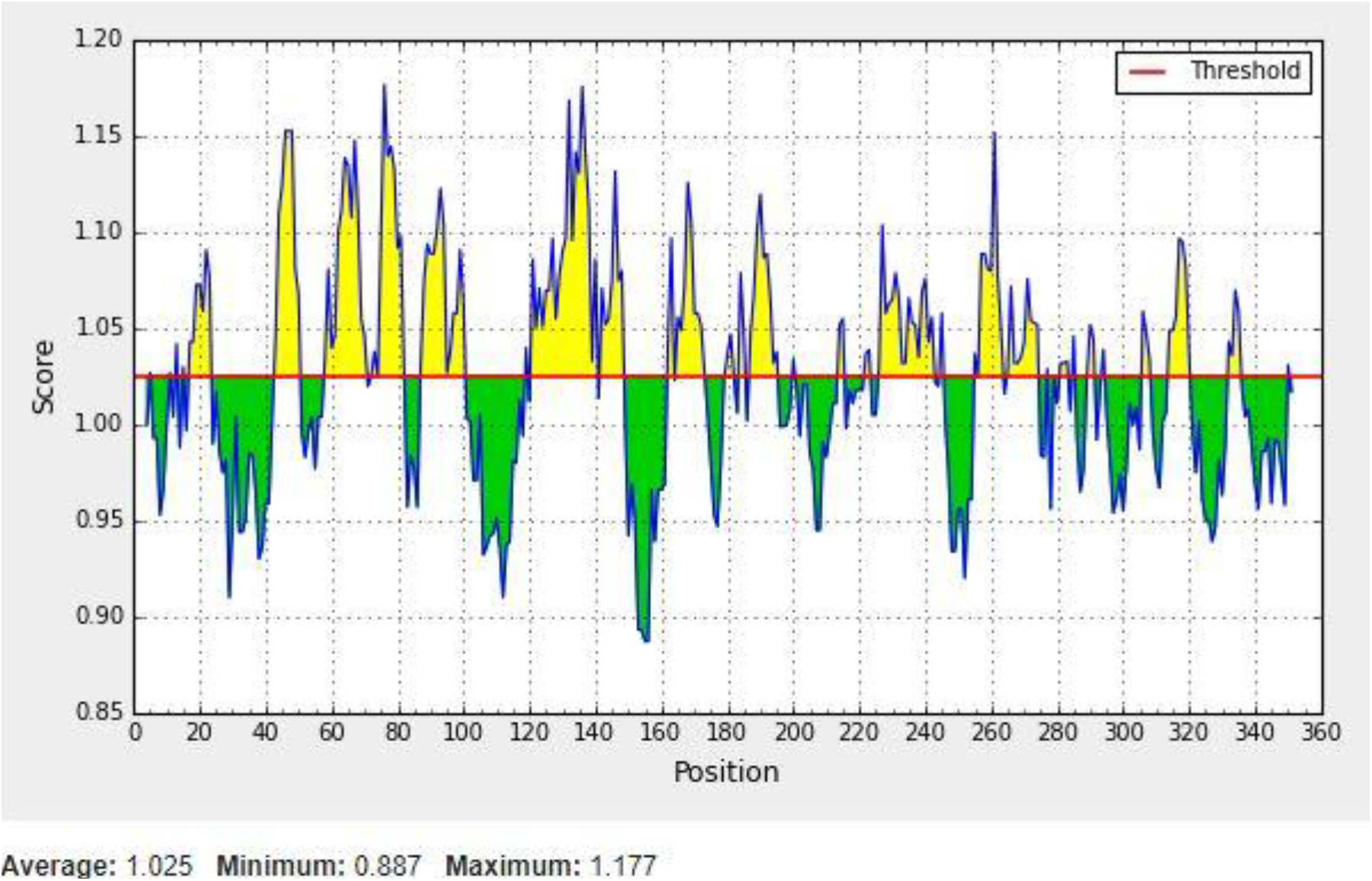
Kolaskar and Tonganokar antigenicity prediction; Yellow areas above the threshold (red line) are proposed to be a part of B cell epitopes and green areas are not.

**Figure 4:**
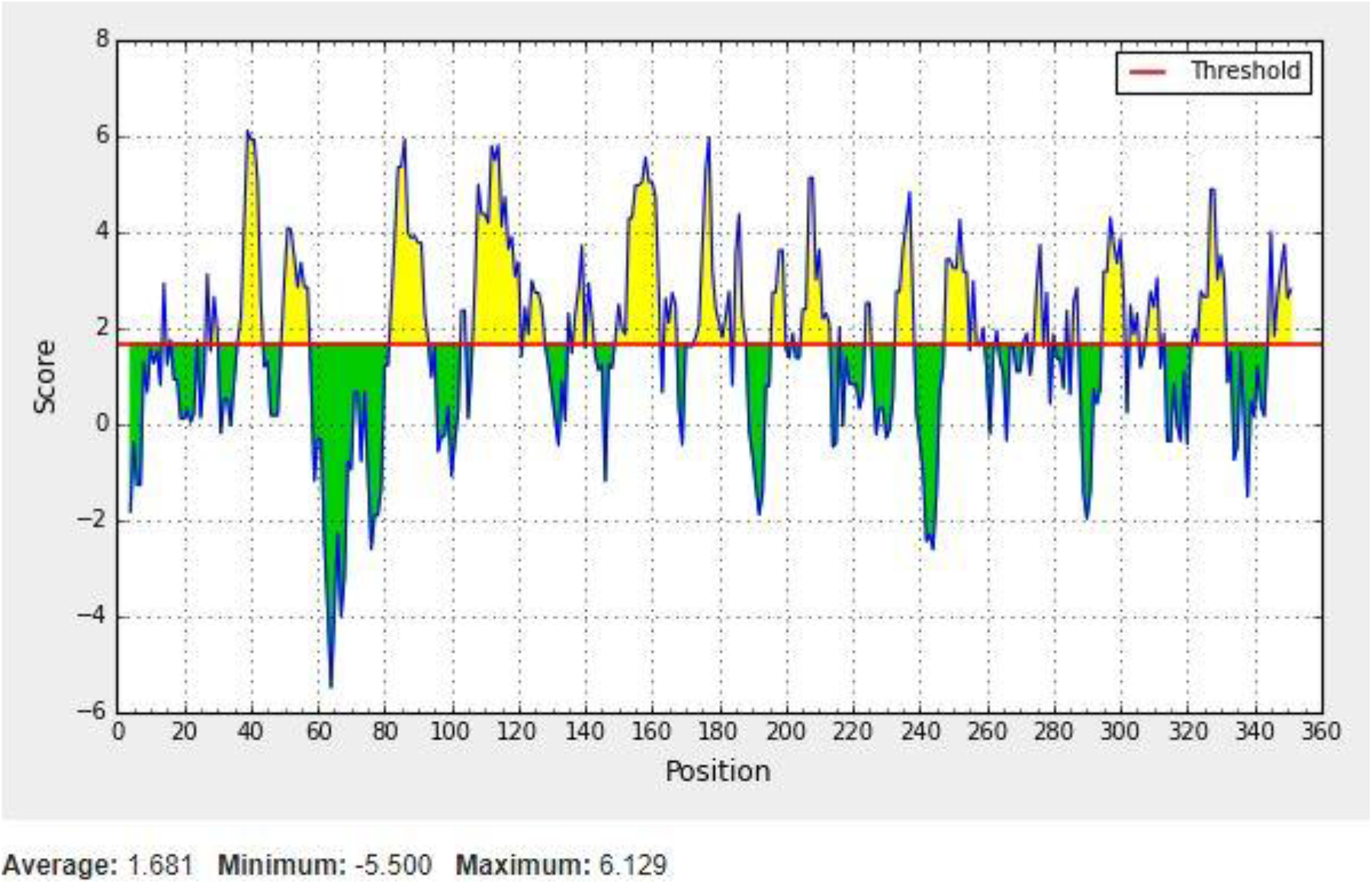
Parker Hydrophilicity prediction; Yellow areas above the threshold (red line) are proposed to be a part of B cell epitopes and green areas are not.

**Figure 5:**
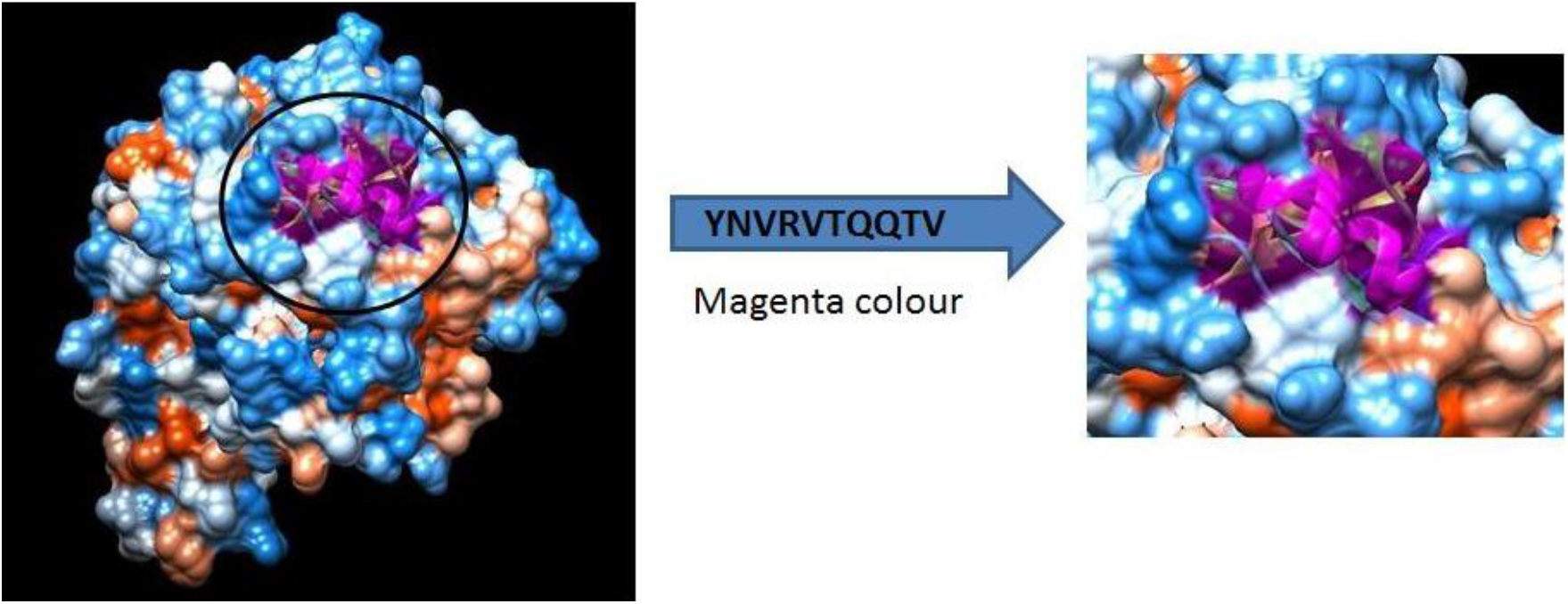
B cell epitopes proposed. The arrow shows position of (YNVRVTQQTV) with Magenta colour in structural level of Fructose 1,6-Bisphosphate Aldolase. *The 3D structure was obtained using USCF Chimera software.

**Figure 6:**
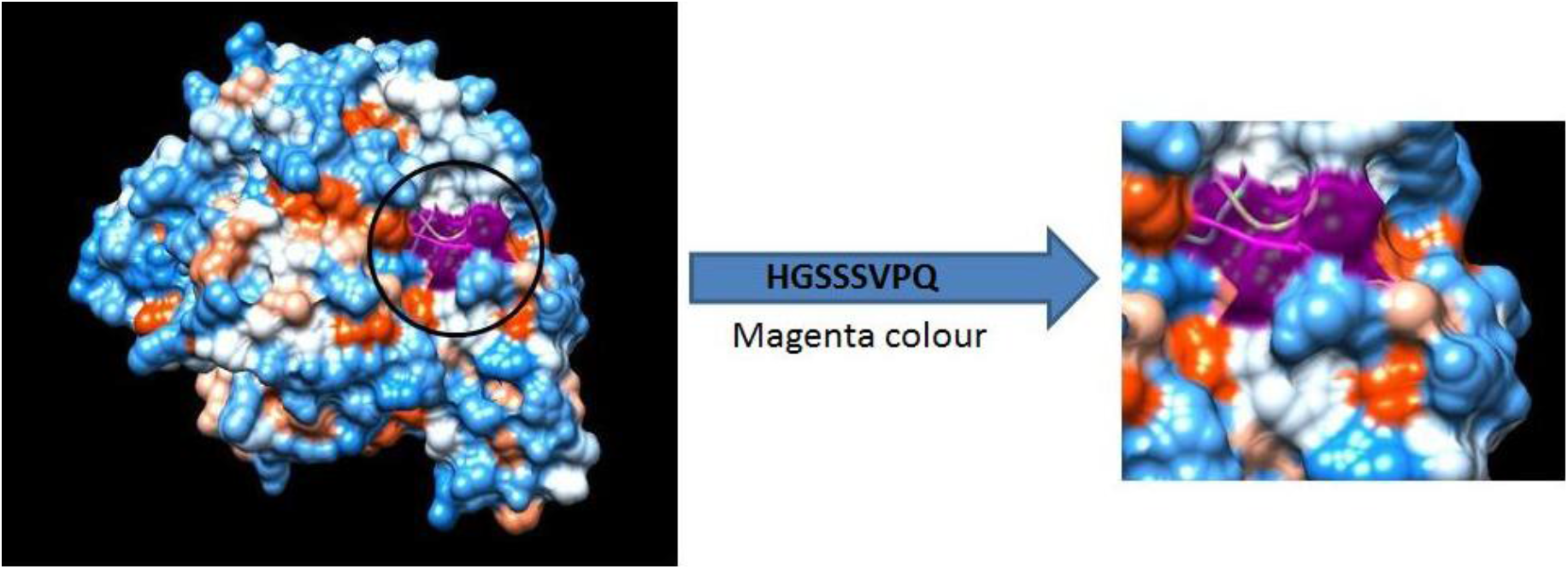
B cell epitopes proposed. The arrow shows position of (HGSSSVPQ) with Magenta colour in structural level of Fructose 1,6-Bisphosphate Aldolase. *The 3D structure was obtained using USCF Chimera software.

**Figure 7:**
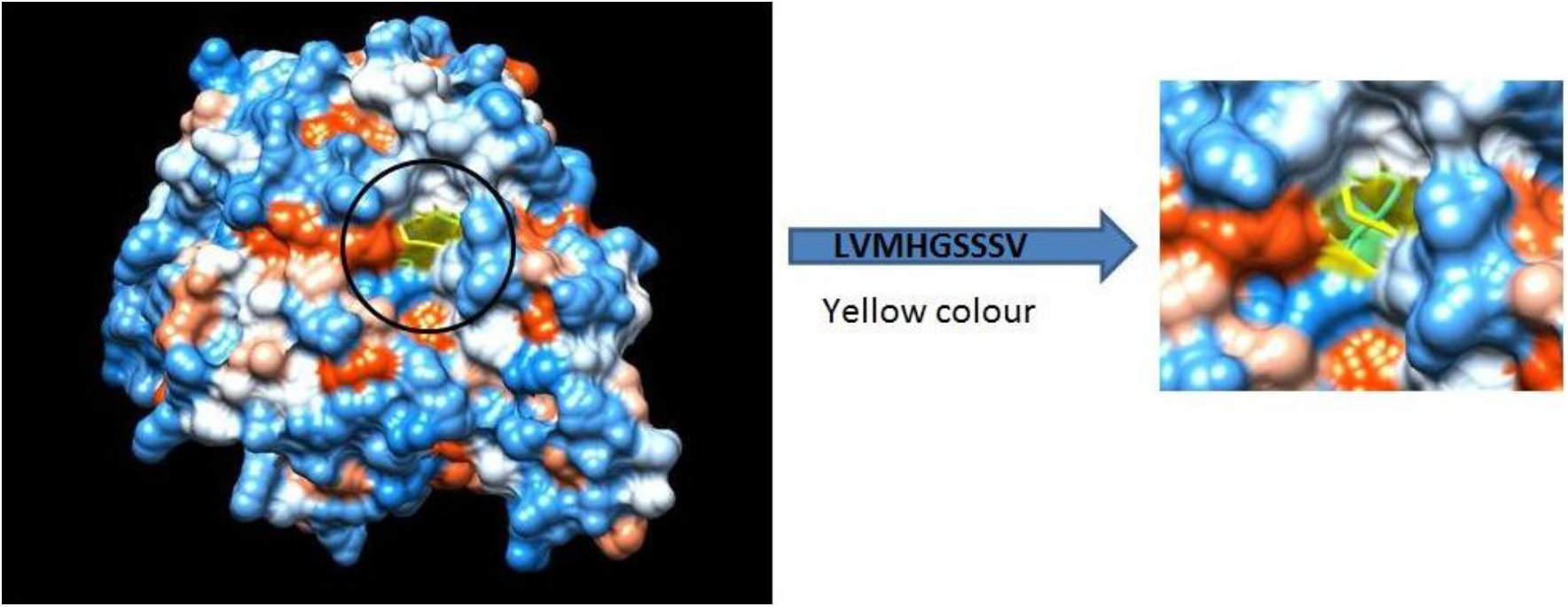
T cell epitopes proposed that interact with MHC1.The arrow shows position of (LVMHGSSSV) with yellow colour in structural level of Fructose 1,6-Bisphopsphate Aldolase. *The 3D structure was obtained using USCF Chimera software.

**Figure 8:**
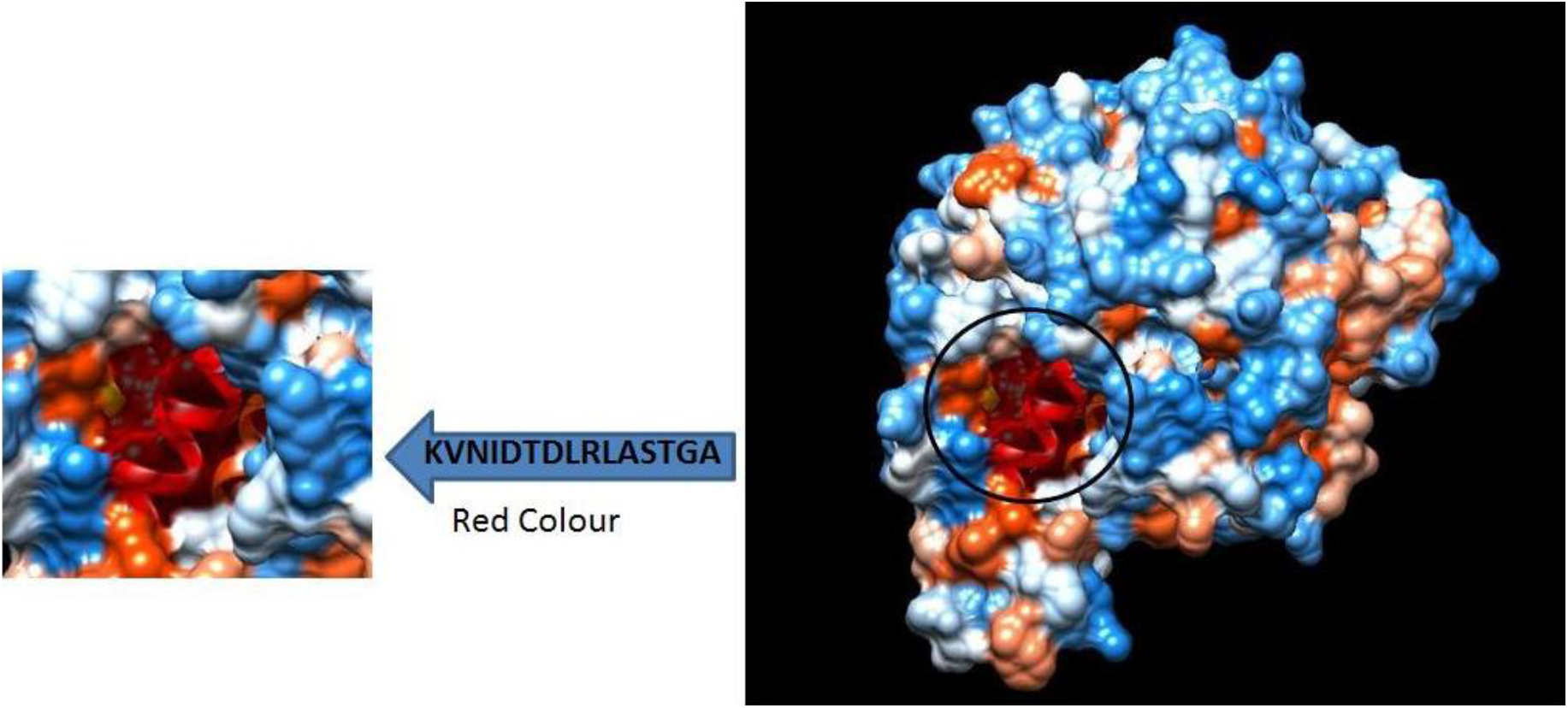
T cell epitopes proposed that interact with MHC2.The arrow shows position of (KVNIDTDLRLASTGA) with Red colour in structural level of Fructose 1,6-Bisphosphate Aldolase. *The 3D structure was obtained using USCF Chimera software.

### Prediction of cytotoxic T-lymphocyte epitopes and interaction with MHC class I

The reference Fructose 1,6-Bisphosphate Aldolase sequence was analyzed using (IEDB) MHC-1 binding prediction tool to predict T cell epitopes suggested interacting with different types of MHC Class I alleles, based on Artificial Neural Network (ANN) with half-maximal inhibitory concentration (IC50) <500 nm. 206 peptides were predicted to interact with different MHC-1alleles.

The most promising epitopes and their corresponding MHC-1 alleles are shown in (Table 2).

**Table 2:**
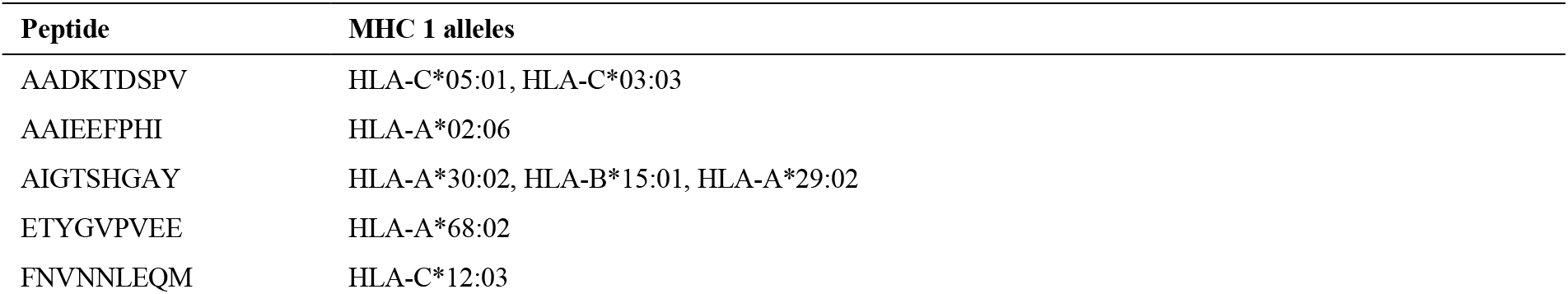
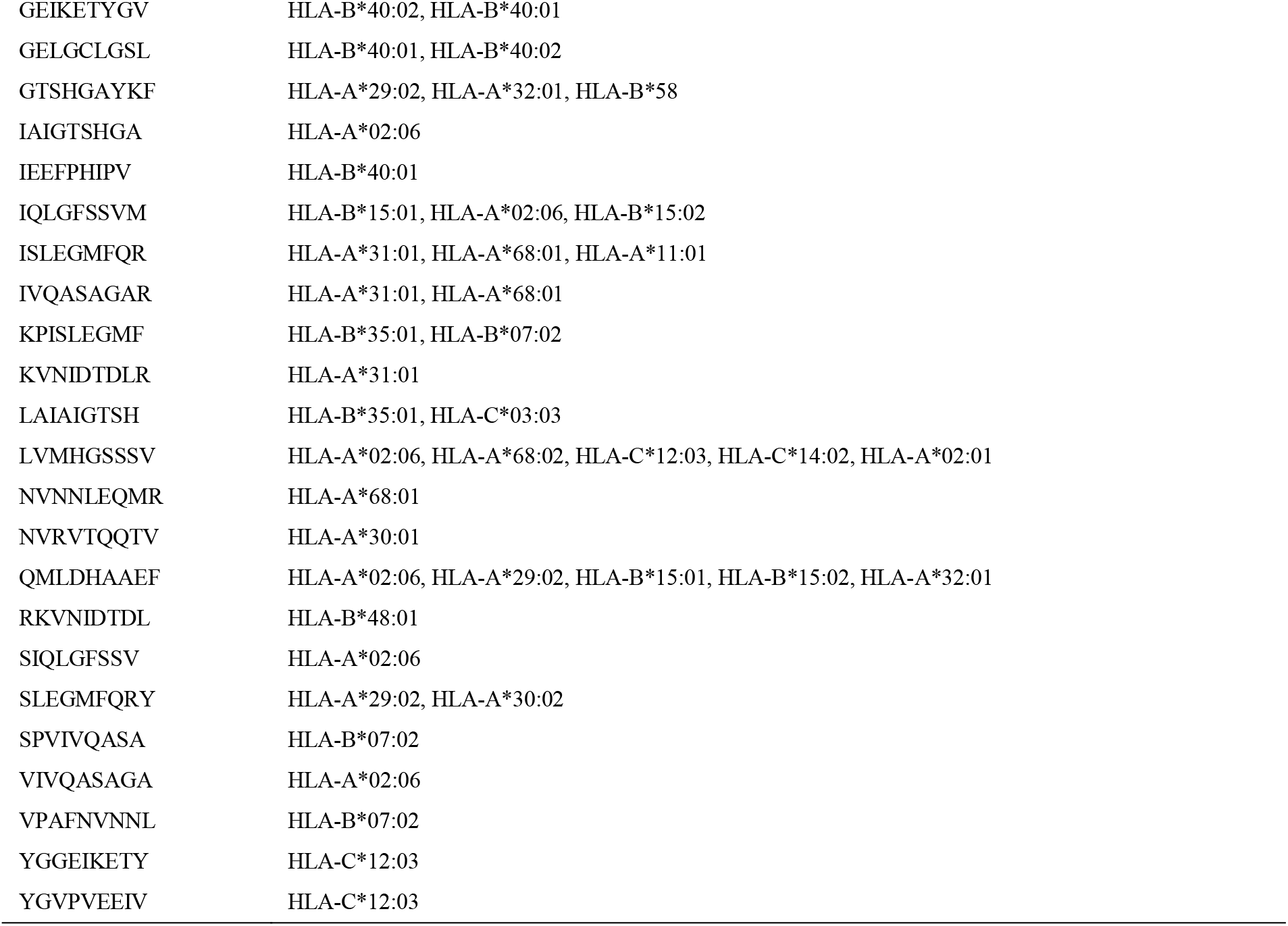
The most promising T cell epitopes and their corresponding MHC-1 alleles.

### Prediction of the T cell epitopes and interaction with MHC class II

Reference Fructose 1,6-Bisphosphate Aldolase sequence was analyzed using (IEDB) MHC-II binding prediction tool based on NN-align with half-maximal inhibitory concentration (IC50) <100 nm; there were 662 predicted epitopes found to interact with MHC-II alleles. The most promising epitopes and their corresponding alleles are shown in (Table 3).

**Table 3:**
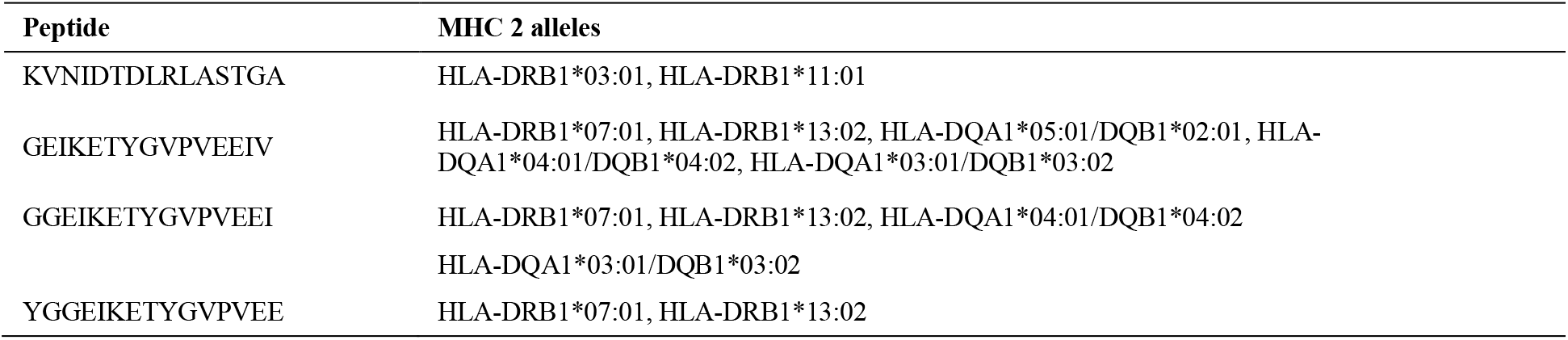
The most promising T cell epitopes and their corresponding MHC-2 alleles.

**Table 4:** The population coverage of whole world for the most promising epitopes of MHC I, MHC II and MHC I & II combined. *In population coverage analysis of MHC II; **8** alleles were not included in the calculation, therefore the above (*) percentages are for epitope sets excluding these alleles: **HLA-DQA1*05:01/DQB1*03:01, HLA-DQA1*01:02/DQB1*06:02, HLA-DQA1*03:01/DQB1*03:02, HLA-DRB4*01:01, HLA-DRB5*01:01, HLA-DQA1*05:01/DQB1*02:01, HLA-DPA1*03:01/DPB1*04:02, HLA-DQA1*04:01/DQB1*04:02**.

| Country | MHC I | MHC II | MHC I,II (combined) |
| --- | --- | --- | --- |
| World | 88.75 % | 61.1 % * | 95.62 % * |

### Population Coverage Analysis

All promising MHC I and MHC II epitopes of Fructose 1,6-Bisphosphate Aldolase were assessed for population coverage against the whole world.

For MHC 1, epitopes with highest population coverage were LVMHGSSSV (60.41%) and QMLDHAAEF (31.7%) (Figure 9 and Table 5). For MHC class II, the epitopes that showed highest population coverage were KVNIDTDLRLASTGA (27.37%) and GEIKETYGVPVEEIV, GGEIKETYGVPVEEI & YGGEIKETYGVPVEE (24.27%) (Figure 10 and Table 6). When combined together, the epitopes that showed highest population coverage were LVMHGSSSV (60.41%), QMLDHAAEF (31.7%) and KVNIDTDLRLASTGA (27.37%) (Figure 11 and Table 7).

**Table 5:**
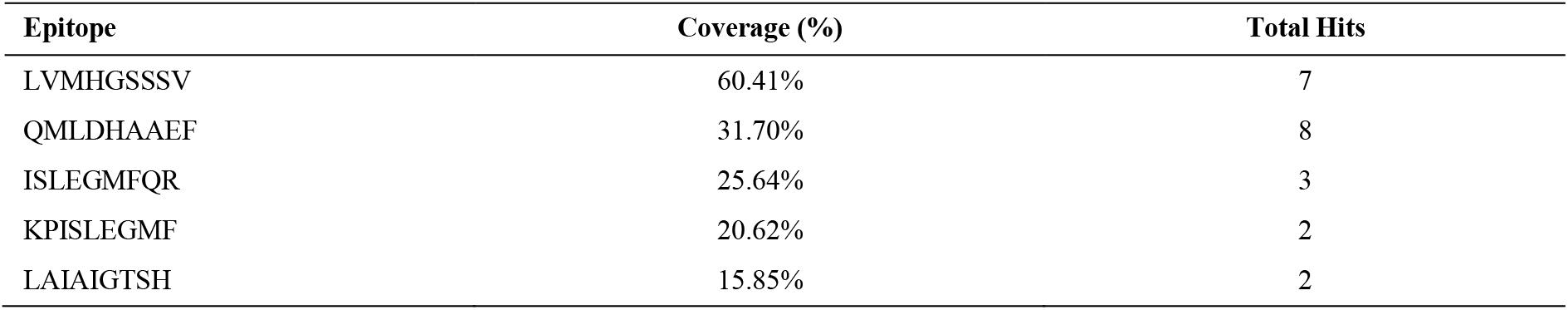
Population coverage of proposed peptides interaction with MHC class I

**Figure 9:**
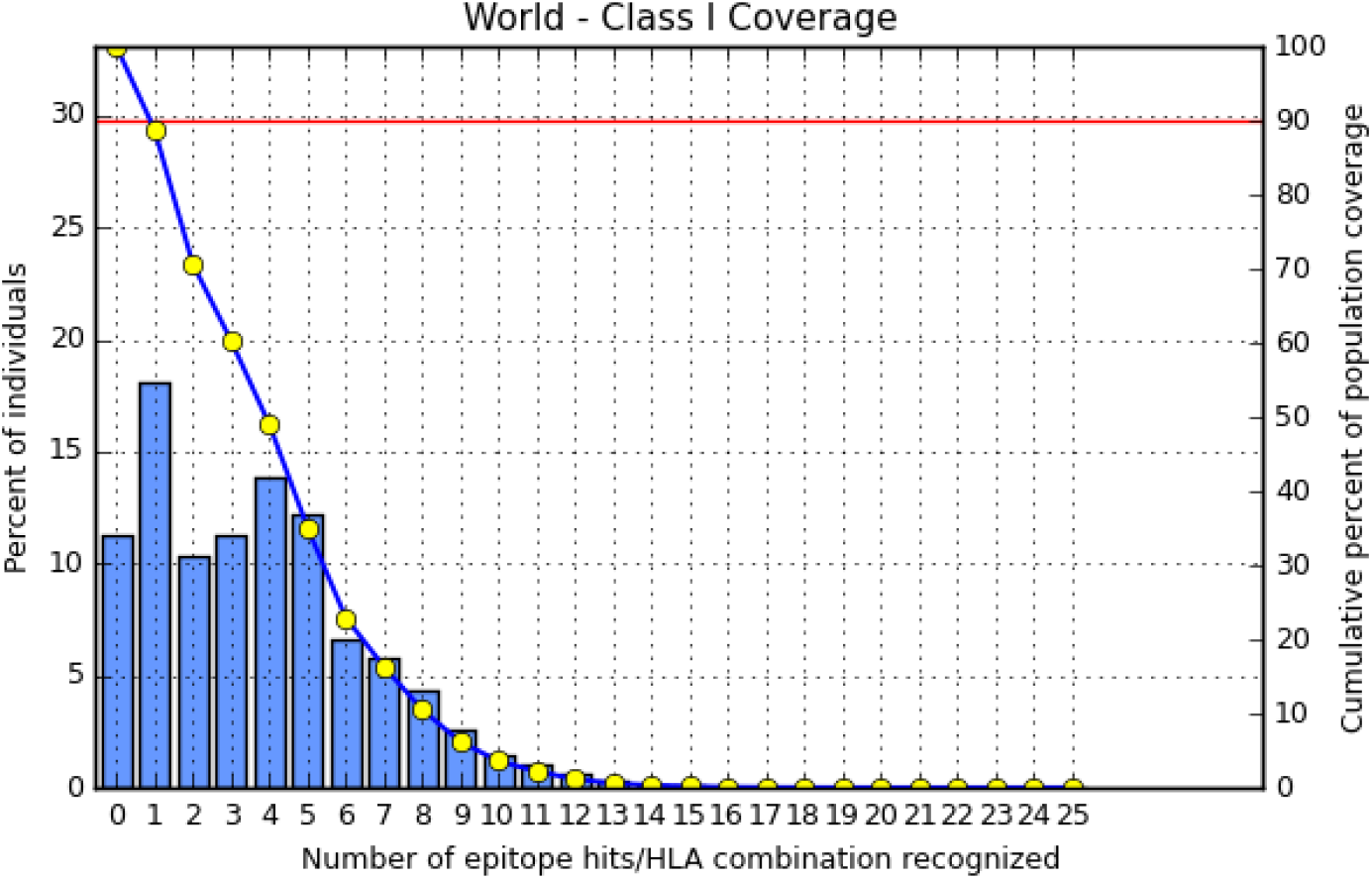
Population coverage for MHC class I epitopes.

**Figure 10:**
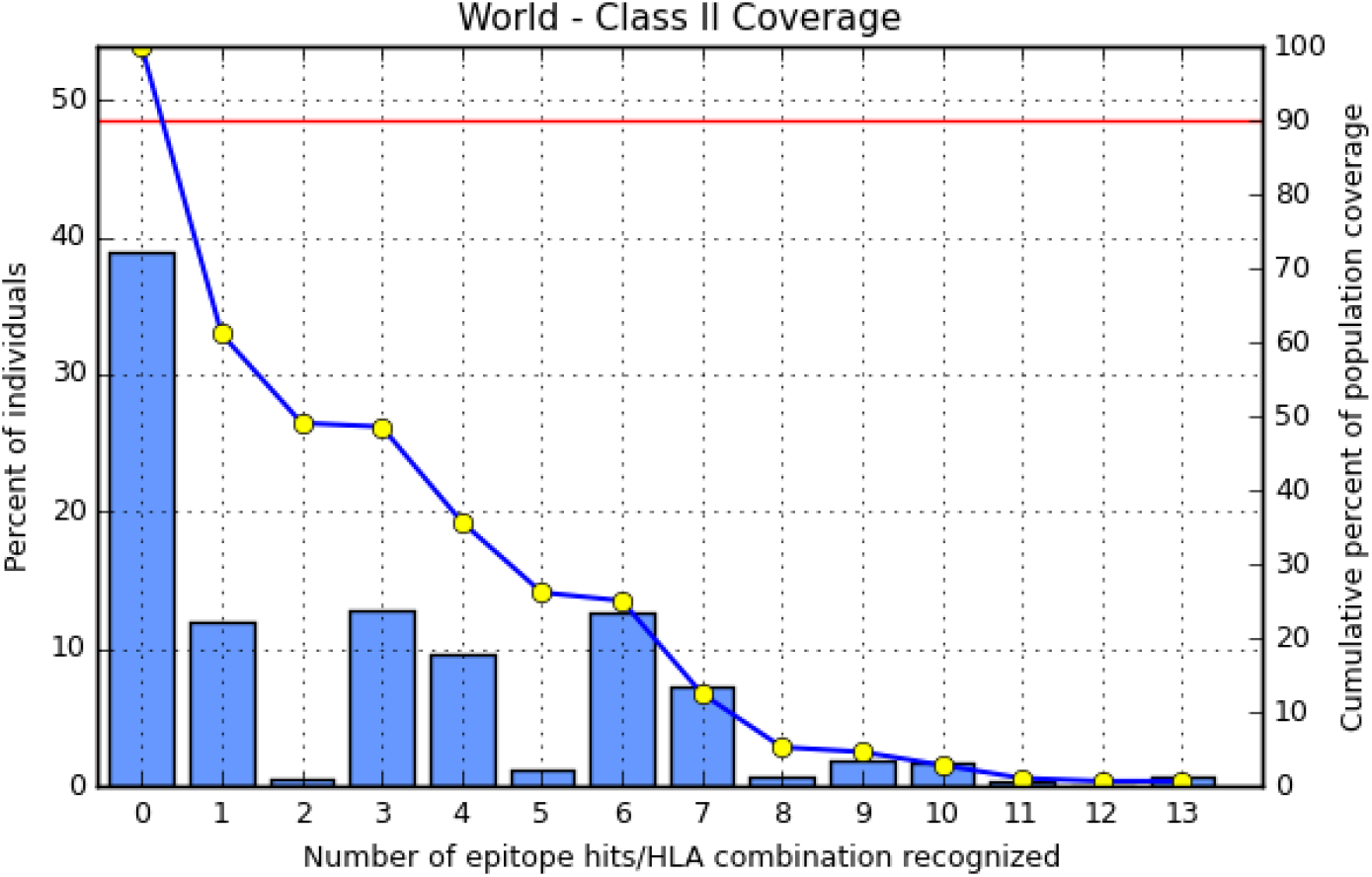
Population coverage for MHC class II epitopes.

**Table 6:** Population coverage of proposed peptides interaction with MHC class 2

| Epitope | Coverage (%) | Total Hits |
| --- | --- | --- |
| KVNIDTDLRLASTGA | 27.37% | 2 |
| GEIKETYGVPVEEIV | 24.27% | 5 |
| GGEIKETYGVPVEEI | 24.27% | 4 |
| YGGEIKETYGVPVEE | 24.27% | 2 |
| GVRKVNIDTDLRLAS | 23.90% | 2 |

**Figure 11:**
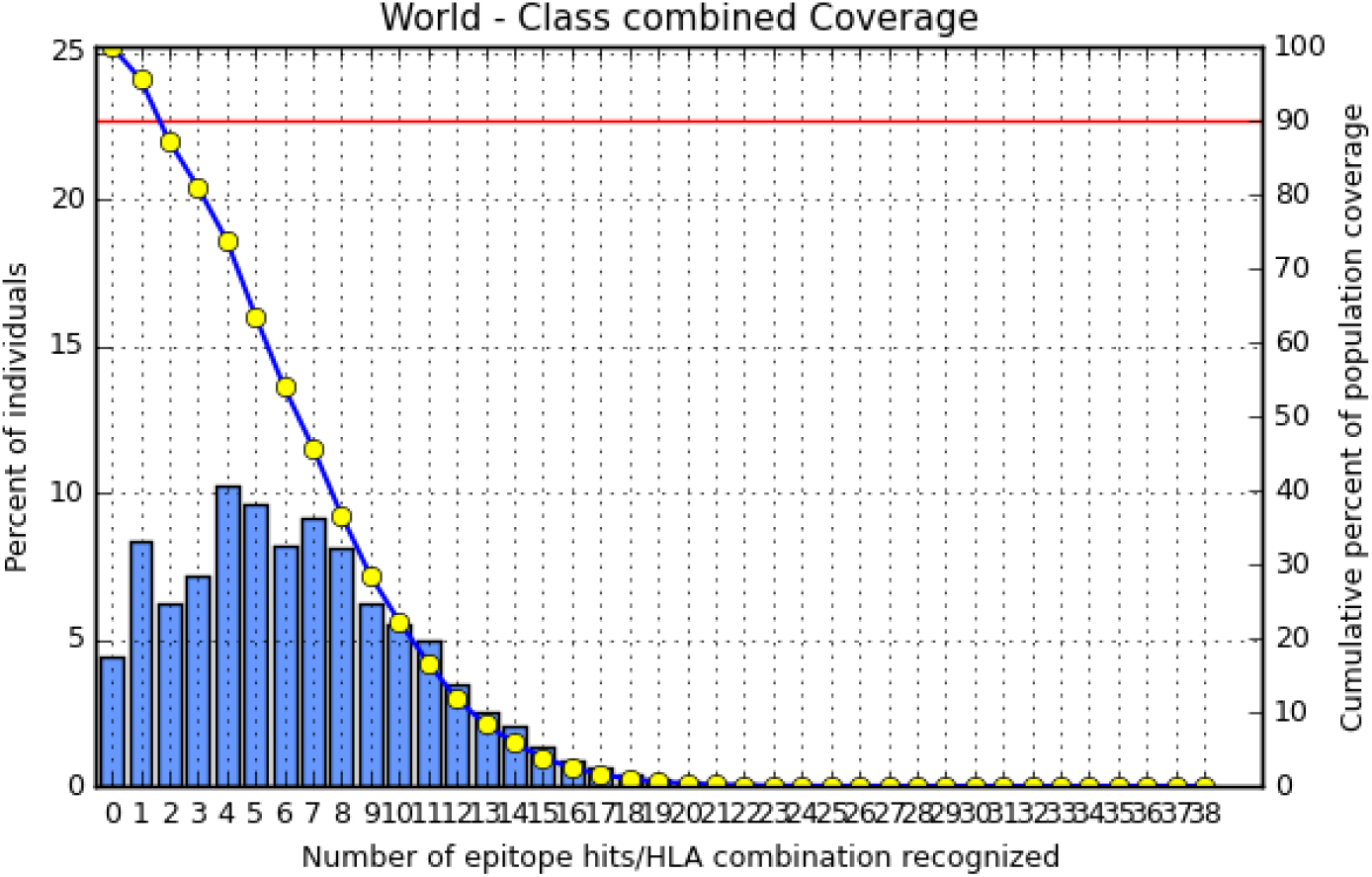
Population coverage for MHC class I & II epitopes combined.

## DISCUSSION

Vaccination against *P. aeruginosa* is highly accredited due to the high mortality rates associated with the pathogen that spreads through healthcare areas. In addition, multidrug resistance of the pathogen demands the design of vaccine as an alternative [43]. In this study, immunoinformatics approaches were used to propose different peptides against FBA of *P. aeruginosa* for the first time. These peptides can be recognized by B cell and T cell to produce antibodies. Peptide vaccines overcome the side effects of conventional vaccines through easy production, effective stimulation of immune response, less allergic and no potential infection possibilities [35]. Thus the combination of humoural and cellular immunity is more promising at clearing bacterial infections than humoural or cellular immunity alone.

As B cells play a critical role in adaptive immunity, the reference sequence of *P. aeruginosa* FBA was subjected to Bepipred linear epitope prediction 2 test to determine the binding to B cell, Emini surface accessibility test to test the surface accessibility, Kolaskar and Tongaonkar antigenicity test for antigenicity, and Parker hydrophilicity test for the hydrophilicity of the B cell epitope.

Out of the thirteen predicted epitopes using Bepipred 2 test, only three epitopes passed the other three tests (ADKTDSPVI, YNVRVTQQTV, HGSSSVPQ) after segmentation. Bepipred version 2 test was used because it implements random forest and therefore predicts large epitope segments.

The reference sequence was analyzed using IEDB MHC1&2 binding prediction tools to predict T cell epitopes. 28 epitopes were predicted to interact with MHC I alleles with half-maximal inhibitory concentration (IC50) < 500. Six of them were most promising and had the affinity to bind the highest number of MHC1 alleles (LVMHGSSSV, QMLDHAAEF, AIGTSHGAY, GTSHGAYKF, IQLGFSSVM, ISLEGMFQR). 19 predicted epitopes were interacted with MHC II alleles with IC50 < 100. Four of them were most promising and had the affinity to bind to the highest number of MHC II alleles (GEIKETYGVPVEEIV, GGEIKETYGVPVEEI, KVNIDTDLRLASTGA, YGGEIKETYGVPVEE). Nineteen epitopes (NVNNLEQMR, IQLGFSSVM, AADKTDSPV, SIQLGFSSV, GEIKETYGV, AIGTSHGAY, VPAFNVNNL, KVNIDTDLR, LAIAIGTSH, IVQASAGAR, ETYGVPVEE, GTSHGAYKF, YGGEIKETY, VIVQASAGA, IAIGTSHGA, RKVNIDTDL, FNVNNLEQM, YGVPVEEIV, SPVIVQASA) appeared in both MHC I and II results.

The best epitope with the highest population coverage for MHC I was LVMHGSSSV (60.41%) with seven HLA hits, and the coverage of population set for whole MHC I epitopes was 88.75%. Excluding certain alleles for MHC II, the best epitope was KVNIDTDLRLASTGA scoring 27.37% with two HLA hits, followed by GEIKETYGVPVEEIV scoring 24.27% with five HLA hits. The population coverage was 61.1% for the all conserved MHC II epitopes. These epitopes has the ability induce T-cell immune response when interacting strongly with MHC I & MHC II alleles effectively generating cellular and humoural immune response against the invading pathogen. When combined, the epitope LVMHGSSSV had the highest population coverage percent 60.41% with seven HLA hits for both MHC I and MHC II.

Many studies had predicted peptide vaccines for different microorganisms such as, Rubella, Ebola, Dengue, Zika, HPV, Lagos rabies virus, and mycetoma using immunoinformatics tools. [44–53] Limitations include the exclusion of certain HLA alleles for the MHC II.

We hope that the whole world will benefit from this epitope-based vaccine and recommend invivo and invitro studies to prove it’s effectiveness.

## CONCLUSION

Vaccination is used to protect and minimize the possibility of infection leading to an increased life expectancy. Design of vaccines using immunoinformatics prediction methods is highly appreciated due to the significant reduction in cost, time, effort and resources. Epitope-based vaccines is expected to be more immunogenic and less allergic than traditional biochemical vaccines.

